# The extinction and survival of sharks across the end-Cretaceous mass extinction

**DOI:** 10.1101/2021.01.20.427414

**Authors:** Mohamad Bazzi, Nicolás E. Campione, Per E. Ahlberg, Henning Blom, Benjamin P. Kear

## Abstract

Sharks (Selachimorpha) are iconic marine predators that have survived multiple mass extinctions over geologic time. Their fossil record is represented by an abundance of teeth, which traditionally formed the basis for reconstructing large-scale diversity changes among different selachimorph clades. By contrast, corresponding patterns in shark ecology, as measured through morphological disparity, have received comparatively limited analytical attention. Here, we use a geometric morphometric approach to comprehensively examine the dental morphology of multiple shark lineages traversing the catastrophic end-Cretaceous mass extinction — this event terminated the Mesozoic Era 66 million years ago. Our results show that selachimorphs maintained virtually static levels of dental disparity in most of their constituent clades during the Cretaceous/Paleogene transition. Nevertheless, selective extinctions did impact on apex predator lineages characterized by triangular blade-like teeth, and in particular, lamniforms including the dominant Cretaceous anacoracids. Other groups, such as, triakid carcharhiniforms, squalids, and hexanchids, were seemingly unaffected. Finally, while some lamniform lineages experienced morphological depletion, others underwent a post-extinction disparity increase, especially odontaspidids, which are typified by narrow-cusped teeth adapted for feeding on fishes. This disparity shift coincides with the early Paleogene radiation of teleosts, a possible prey source, as well as the geographic relocation of shark disparity ‘hotspots’, perhaps indicating a regionally disjunct pattern of extinction recovery. Ultimately, our study reveals a complex morphological response to the end-Cretaceous mass extinction event, the dynamics of which we are only just beginning to understand.

## INTRODUCTION

Fossils provide the only direct evidence for the interplay between organisms and their environments over vast evolutionary timescales [1–3]. They are, therefore, crucial for exploring the drivers of biodiversity and ecosystem change in the past, with potential insights on the origin of present-day ecosystems [3]. However, the analytical challenge is to discern a genuine biological signal relative to geologic, taphonomic, sampling, and philosophical biases [4–7]. While these may be impossible to overcome in entirety, the fossil records of some widely distributed and chronostratigraphically extended clades offer reasonable proxies for modelling macroevolutionary processes through deep time.

Sharks constitute just such a ‘model group’ because their dental remains are abundant in Mesozoic and Cenozoic marine deposits — a timeframe that covers ∼250 million years (Ma) [8,9]. Extant shark species are also ecologically ubiquitous, encompassing a spectrum of macrophagous to microphagous predators that account for nearly half (42%) of all the currently documented chondrichthyan biodiversity (n=1193 species) [10,11]. Nevertheless, after nearly 200 years of scientific research [9], the various biological and environmental factors that shaped shark evolution in the distant past remain obscure. In particular, their capacity to survive past mass extinctions is relevant for understanding the dramatic decline of shark populations observed in modern oceans [11–15].

The end-Cretaceous mass extinction (∼66 Ma), which marks Cretaceous/Paleogene (K/Pg) chronostratigraphic boundary, is especially pertinent because it profoundly disrupted marine ecosystems, but has had disputed implications for shark species diversity and morphological disparity. Indeed, contrasting interpretations have advocated either limited [16], or complex interrelationships between various biotic and abiotic drivers influencing shark evolution from before, during, and after the K/Pg mass extinction event [17–20]. Here, we attempt to resolve the extinction dynamics and inferred mechanisms of sharks via a comprehensive assessment of their dental morphological disparity across the end-Cretaceous mass extinction. Our approach expands on previous studies that have targeted either geographically localized [21,22], or clade-specific [20] assemblages. We use a dataset of 1,239 fossil shark teeth, representing nine major selachimorph clades sampled at global and regional scales. These groups include: the Galeomorphii orders Carcharhiniformes, Heterodontiformes, Lamniformes, Orectolobiformes; Squalomorphii orders Echinorhiniformes, Hexanchiformes, Squaliformes, Squatiniformes; and the extinct [†]Synechodontiformes. Our geometric morphometric analysis compares changing patterns of dental disparity and morphospace across a constrained 27.6-million-year interval spanning the Campanian and Maastrichtian ages of the Late Cretaceous (83.6–66 Ma), to the Danian, Selandian, and Thanetian ages (=Paleocene epoch) of the early Paleogene (66–56 Ma). Using this new dataset, we test the following hypotheses via their associated predictions. 1) Long-term shifts in Earth Systems in the twilight of the Mesozoic, including major global regressions during the Maastrichtian, saw to a loss of marine habitats [22,23]. We, therefore, predict that shark diversity and disparity were in decline prior to the mass extinction event. 2) The end-Cretaceous was a devastating period for sharks, as hypothesized by major losses in the diversity of species, genera, and families [17,18]. Such losses, especially at higher taxonomic ranks, suggest concomitant losses in ecological diversity, which should be expressed by notable disparity decreases at the Boundary. 3) Previous studies hypothesized a non-random, selective extinction of sharks, in particular against pelagic large-bodied pelagic apex predators and shallow-water carpet shark [17,20,24]. As a result, we predict that the extinction event was biasedly more catastrophic towards specific morphotypes (e.g., ‘cutting-type’ dentitions), while others remained largely unaffected. 4) The end-Cetaceous extinction was a global event and, as such, we predict that all major late Mesozoic ocean habitats were consistently affected, leading to similar disparity profiles at both regional and global scales. 5) The aftermath of the extinction saw to a major radiation of teleost fish [16], a potentially important food source for the surviving sharks. As such, we predict a Paleocene proliferation of morphologies associated with bony-fish diets, such as elongate, thin teeth of the ‘piercing’ and ‘tearing-type’ designs.

## MATERIALS AND METHODS

### Dataset assembly

Our dataset of 1,239 teeth includes photographs or images of fossil selachimorph teeth taken from first-hand observations or compiled from the literature (Appendix S1; S1 Fig and S1 and S2 Table). Following recommended best-practices [20,25,26], our raw image data was screened to only include tooth specimens with complete crowns and of adequate resolution, so as to unambiguously determine the position of the root-crown junction. Our global coverage includes sharks from nearly all taxonomic orders (S1 and S2A Fig), with the exception of Pristiophoriformes (Sawsharks), which have a notoriously sparse fossil record [8,27]. We also elevated Echinorhinidae to Echinorhiniformes [28–30], and employed a sensitivity analysis to test the morphospace occupation and disparity effects of †Synechodontiformes, which are classified as either a clade of Galeomorphii, or a neoselachian sister lineage (S2A Fig) [31–34]. Finally, all images were standardised so that the tooth apex was directed to the left and were generally digitized in the preferred labial aspect, unless only a lingual view was figured (S3 Fig). The equivalency of labial and lingual views was demonstrated in [20] but we nonetheless verify it again here, through the use of ordinary least-squares linear models and a sub-sample of the data for which both labial and lingual views were digitized.

Our dataset was time-binned across all five geochronological ages constituting the immediate K/Pg interval: Campanian + Maastrichtian/Danian + Selandian + Thanetian. However, we also implemented an alternative four-age time-binning scheme that pooled temporally ambiguous specimens assigned to the Danian + Selandian (S3 and S4 Table). In addition, we carried out analyses using a subsample of the dataset (N=829) for which sub-ages (early, middle, and late) of the Campanian, Maastrichtian, and Danian could be defined (S5 Table). Global taxonomic richness (henceforth referred to as just diversity) at genus-level and morphological disparity were plotted against the mean Ma values taken from the International Chronostratigraphic Chart v2020/03 [35].

### Acquisition of geometric shape data

Landmark-based geometric morphometrics quantifies biological shapes as a series of evolutionary homologous points in Cartesian space [25,26,36–41]. In sharks, however, the evolutionary homology [37,42] between landmarks cannot be assumed due the inherent morphological variability of shark teeth (e.g., the location and number of cusplets). As a result, the landmark types and placements are argued on the basis of topological homology [40].

Landmark digitization was carried out in *tpsDig2* v. 2.31 [43] with resampling to a standard number of equidistant semilandmarks carried out using customized code in *R* v. 3.6.3 [44]. The resulting landmarking scheme comprise two open curves defined by semilandmarks and anchored by three fixed landmarks: two Type 1 landmarks delimited the mesial and distal crown-root junctions and one Type 2 landmark associated with the tooth apex (sensu [42]; S2B Fig and S6 Table). To identify the desired number of semilandmarks (k), the mesial and distal curves were resampled seven times to increasing numbers of equally spaced semilandmarks: k = 40, 60, 80, 100, 120, 140, and 160. Qualitative observations (S4 and S5 Fig) found that k = 160 best-captured tooth shape complexity (e.g., adequate shape coverage of hexanchiforms) via distal and mesial curves of 78 and 79 sliding semilandmarks, respectively. At this time, tooth serrations were not digitized due to image resolution, but we acknowledge that they are functionally important [45].

Finally, to screen for possible digitization errors, we extracted a random sub-sample of 30 images (S7 Table) and used a one-way ANOVA to calculate the intra-class correlation coefficient (*R*) [25,46,47] (also see Supplementary text). We also ran a two-block partial least-squares (2B-PLS) analysis to infer covariation between the datasets. Other exploratory procedures performed as part of assessing measurement error include: 1) a manual survey of digitized images to confirm accurate landmark placement and 2) screening for outliers. The latter was performed visually, via observation of ordinated shape-space and associated thin-plate spline (TPS) deformation grids and analytically, through an outlier search function, *plotOutliers*, in the geomorph package *geomorph* v. 3.3.1 [48].

### Morphometric analysis

To standardize digitized specimens to unit size, position, and rotation, we use a generalized Procrustes analysis (GPA) [49,50] that minimizes the bending energies to optimize the positions of the sliding semilandmarks [50,51]. Because large numbers of sliding semilandmarks can impinge on GPA to reach proper convergence [25,37], we varied the iteration frequency by arbitrarily increasing the *max*.*iter* argument in *gpagan* to compare convergence criteria Q-values (= Procrustes sum of squares) and then inspected the resulting consensus shape configurations (S6 Fig and S8 Table).

The aligned Procrustes coordinates were ordinated via a principal components analysis (PCA) based on the singular value decomposition of the variance-covariance matrix. Shape variation was depicted as both TPS deformation grids and deformation isolines to generate a concentration ‘heat map’ [52]. Ordinated space was visualised via back-transformation [53,54]. All analyses were carried out in *R* v. 3.6.3 [44] with the *geomorph* v. 3.3.1 [48] and *RRPP* v. 0.6.2 [55] packages; visualization used the *ggplot2* package [56]. All data and *R* scripts are available from *Data Dryad*.

### Temporal analyses of morphospace

Ordinated morphospace was categorized as time-bin boxplots that incorporated the arithmetic mean, median, and modal values. Confidence intervals were calculated using non-parametric bootstrapping with 1000 resamples. Comparisons between multiple central tendency values accommodated for differences in frequency distribution. Modal shape configurations corresponded to the region of maximum frequency calculated as:

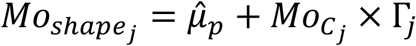

where 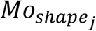 is the modal shape configuration along the *j*th PC; 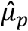 represents the mean shape configuration of the whole sample; 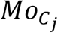 is the mode of the *j*th PC axis; and, Γ_j_ is the rotation matrix corresponding to the *j*th PC. 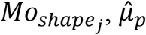 and Γ_j_ take the form of k × m matrices and were plotted as TPS deformation grids; k is the number of landmarks and m is the dimensions, in our case 2.

Non-normal distributions were assessed using a Henze-Zirkler’s test [57] for multivariate normality (*HZ*=4956, *p*=0) (S7 Fig). Statistical comparisons between time-bins used a non-parametric Procrustes analysis of variance (PAV) implemented in the *RRPP* package [55].

### Temporal analyses of disparity

We used Procrustes variance (PV) to calculate disparity [40,58] from our 2D (k × m × N) landmark dataset as:

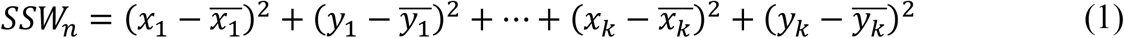

where *SSW*_*n*_ is the sum of the square distances between the coordinates (*x*_*k*_ and *y*_*k*_) of observation *n*, and their associated mean (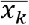 and 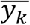)This may be alternatively depicted as:

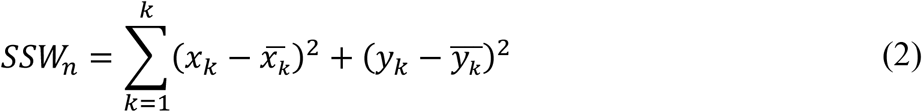

All *SSW*_*n*_ values in a given time-bin *t* are then summed and divided by the sample size at that time (*N*_*t*_) to measure PV, across all observations.

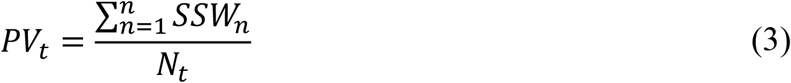

Disparity within each time-bin was partitioned according to our taxonomic order-level classifications, which determined the clade-specific contributions to overall disparity. This calculation equates to (2), but with distances measured relative to the mean of the group 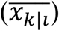, rather than the overall mean [48]:

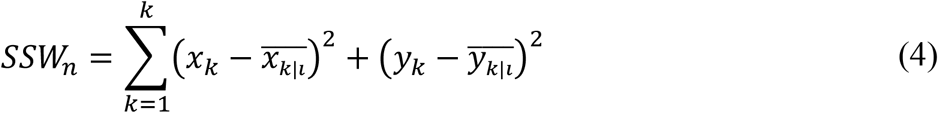

The partial *PV* for the group *i* over a specific time-bin *t* is ∴

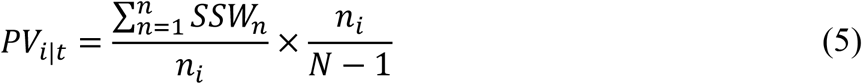

where *n*_*i*_ is the sample size within group *i* and *N* is the total sample size within *t*.

Computationally, these equations are solved in *geomorph* [48], with the expectation that additive partial disparities [59] for sampling within each time-bin approximate the total *PV* given *t*.

We used a residual randomization permutation procedure (RRPP) with 1000 permutations [55] to test null hypotheses for our multivariate shape data as:

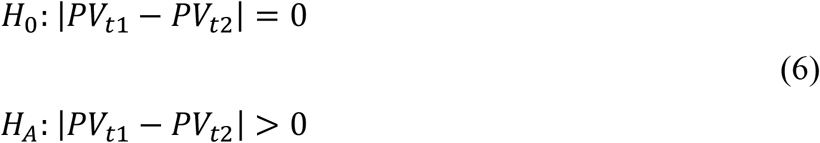

where *H*_*0*_ assumes that pairwise absolute differences between PVs across two given time-bins (e.g. *t1* and *t2*) will be zero. *H*_*A*_ alternatively stipulates that the difference will be greater than zero.

We also applied non-parametric bootstrap resampling to estimate confidence intervals around disparity. All *post-hoc* pairwise comparisons of group-means were subject to false discovery rate (FDR) adjustments for *P*-values to mitigate the increased risk of Type I errors associated with multiple comparisons [60].

### Geographic distribution in the fossil record

We accommodated for inherent sample size biases in the fossil record [4,61–63] via rarefaction to aid comparisons of time-scaled PV [20]. This involved sub-sampling all time-bins to a minimum time-bin size (S1, S3 and S4 Table) 999 times, from which 95% prediction intervals were calculated. Geographic sub-sampling focused on the UNESCO World Heritage fossil locality at Stevns Klint in Denmark (S8 Fig), which preserves exceptionally rich selachimorph assemblages [64,65] spanning the K/Pg succession [66]. However, we also calculated partial disparities for each time-bin based first on depositional basins, and then using countries of origin; designated *i* in (5). Although geopolitical boundaries are artificial, they provide a convenient proxy for comparing regional versus global disparity signals across a broader sub-sample series.

### Influences of heterodonty

Sharks are known to exhibit both monognathic (MH, variation along the tooth row) and dignathic (DH, variation between the upper and lower jaws) heterodonty [8,67], which is usually undetectable with isolated fossil teeth [68]. To attempt to remove potential confounding factors associated with tooth position that may affect morphospace distributions [20,69] we inferred all teeth to either parasymphyseal, anterior, lateroposterior, and posterior tooth positions (N_MH_=897) and segregated teeth from the upper and lower tooth rows (N_DH_=334) (also see Supplementary Text).

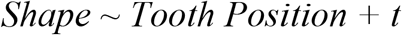

where the aligned Procrustes coordinates (*shape*) are described as a function of monognathic versus dignathic tooth position and age (*t*).

Lastly, because developmental [69,70] and ontogenetic factors [69,71–73] are likewise difficult to define from isolated teeth, we conducted our analyses with the caveat that adequate intraspecific coverage was assumed for each order-level clade.

### Taxonomic richness

We used an occurrence-based sub-sample to calculate taxonomic richness of sharks at genus-level. Classic rarefaction (CR) and Shareholder Quorum Sub-sampling (SQS) [61,74] were implemented in the *divDyn* v. 0.8.0 package [75]. This employed a four-age time-binning scheme with rarefaction constrained at a sub-sample size of 30, and the SQS quorum level at 0.6 ([76]). Results were visualized as quantile plots.

## RESULTS

### Digitization measurement error

Visual comparison of the computed consensus (mean) tooth shapes (S9 Fig) indicated very low error margins. Accordingly, an intra-class correlation coefficient (*R*) of 2% (or 1 in 50) was calculated based on the aligned Procrustes coordinates and their error replicate counterparts (N=30) (S9 Table), as well as Pearson’s product-moment correlation (t=1874.7, *df*=9598, *p*-value << 0.001, *R*=0.99) and two-block partial least squares tests (r-pls=0.997, *p*-value=0.001, Z=7.136), which unambiguously demonstrate dataset compatibility (S10 Fig).

### PCA visualization

PC1–4 explains 89.28% of the shape variation (Fig 1A–C, S11 Fig), with the other PC axes describing less than 5% (S11 Fig and S10 Table). PC1 (62%) captures tooth height and width variation from apicobasally tall and narrow teeth, to mesiodistally broad and low crowns (Fig 2A). PC2 (12%) alternatively represents distally recurved teeth with low ‘heels’, versus upright triangular teeth with lateral cusplets (Fig 2B). PC3 (11%) tracks tall and conical, to distally wide and recurved teeth with pronounced lateral cusplets (Fig 2C). PC4 (5%) captures a spectrum of short triangular teeth with reduced cusplets, to tall crowns with prominent cusplets that equalled the main cusp in height (Fig 2D).

**FIG. 1.**
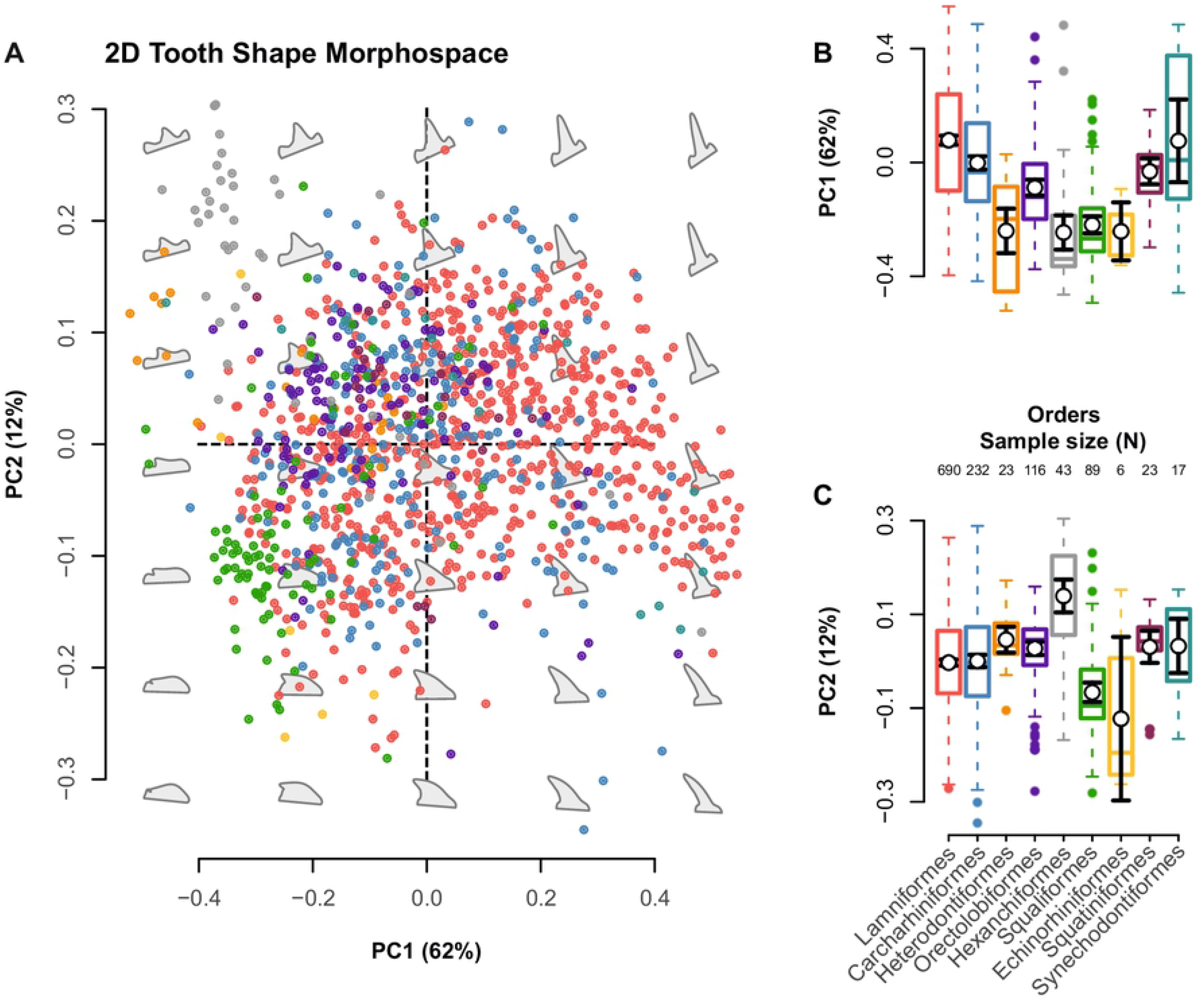
Morphospace distribution. **(A)** Multivariate shape space for the n=1239 global selachimorph teeth sample. Theoretical backtransform tooth shapes (grey) indicate shape variability across morphospace as defined by PC1 and PC2. **(B–C)** Box-and-whisker plots indicating average morphospace occupation along **(B)** PC1, and **(C)** PC2. Error bars represent 95% confidence intervals. Proportion of variance and group sample sizes are listed in the axis labels.

**FIG. 2.**
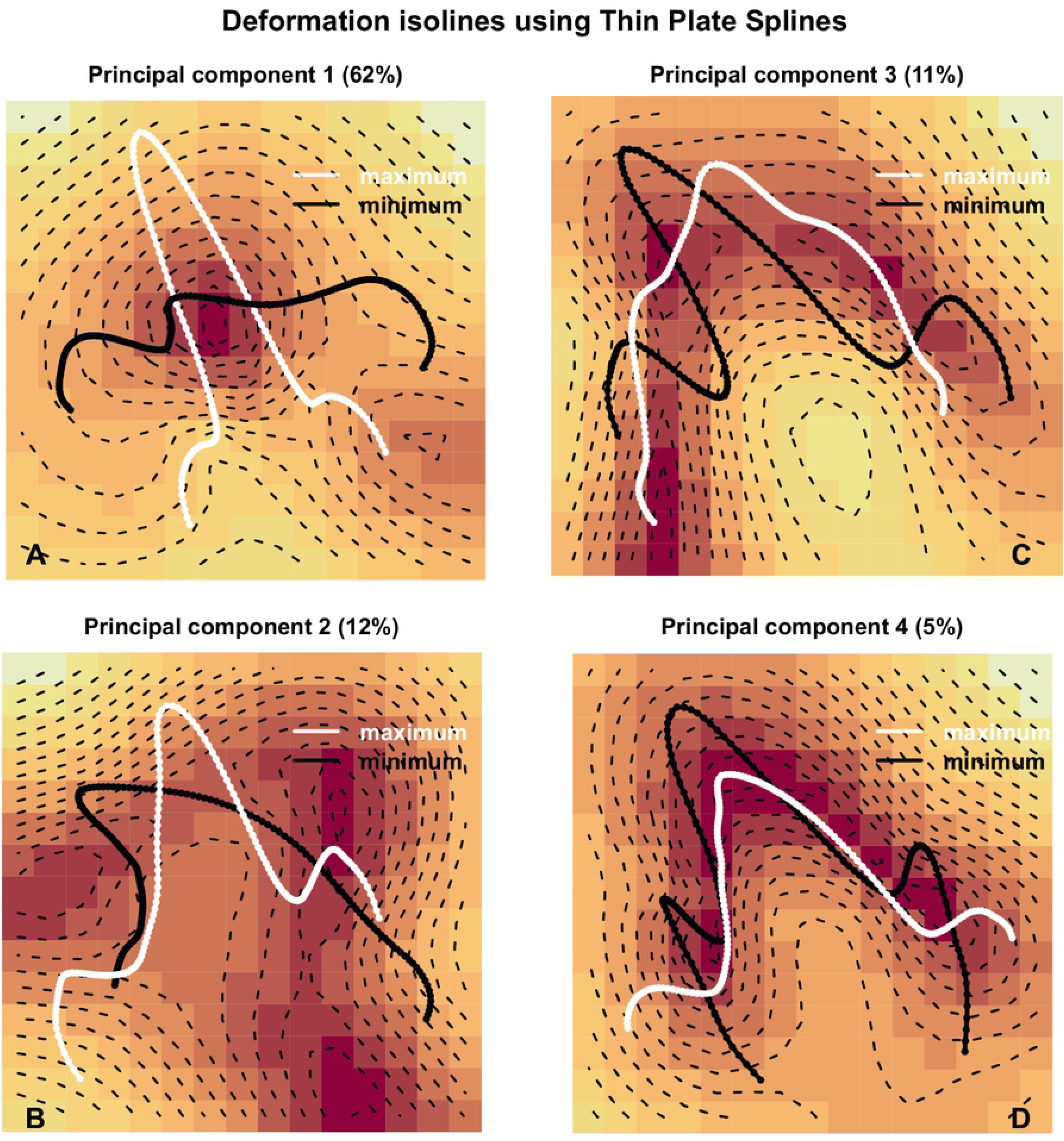
Hypothetical morphologies corresponding to the minimum and maximum values of PC1–PC4. **(A–D)** Isolines and counter lines based on the TPS-method. Areas of concentrated strain are indicated by increasing red intensity.

While there was substantial overlap in tooth morphologies between our sampled selachimorph clades (Fig 1A–C), a general Procrustes one-way ANOVA found significant difference between the group means and specific axes of variation (Table 1). This accords with visual segregation between the clades in morphospace (Fig 1A–C), and is further illustrated by relative measures of central tendency, distribution symmetries, test of normality and multimodality (S11Table).

**Table 1.** Non-parametric analysis of variance based on a randomized residual permutation procedure (RRPP) for. n=1239. Coefficient estimation via ordinary least squares (OLS). Type I (sequential) sums of squares were used to calculate the sums of squares and cross-products matrices. Effect sizes (Z) are based on the F distribution.

|  | *d.f.* | SS | MS | R<sup>2</sup> | F | Z | Pr(>F) |
| --- | --- | --- | --- | --- | --- | --- | --- |
| **All PC-axes** |  |  |  |  |  |  |  |
| ~ groups (clades) | 8 | 18.744 | 2.34294 | 0.18931 | 35.904 | 10.892 | **0.001\*\*** |
| **PC1** |  |  |  |  |  |  |  |
| ~ groups (clades) | 8 | 13.737 | 1.71707 | 0.22474 | 44.569 | 7.3642 | **0.001\*\*** |
| **PC2** |  |  |  |  |  |  |  |
| ~ groups (clades) | 8 | 1.4895 | 0.186190 | 0.12352 | 21.668 | 5.8106 | **0.001\*\*** |
| **PC3** |  |  |  |  |  |  |  |
| ~ groups (clades) | 8 | 2.1710 | 0.271372 | 0.20412 | 39.431 | 7.0403 | **0.001\*\*** |
| **PC4** |  |  |  |  |  |  |  |
| ~ groups (clades) | 8 | 0.3986 | 0.049819 | 0.08708 | 14.665 | 5.1632 | **0.001\*\*** |

Symbols: *d.f.* = degrees of freedom; SS = sequential sums of squares; MS = Mean squares; F statistics = F value
by permutation; R<sup>2</sup> = coefficient of determination; Z = effect size. *P*-values are based on 999 permutations.

### Global and regional disparity

We found overall stability in selachimorph global disparity across the Campanian–Thanetian interval (Fig 3A, S12 Table). The only exception was a significant decline within the Selandian time-bin (PV_Danian_=0.082; PV_Selandian_=0.042; *p*=0.033), which may be a product of sampling (N=28) and/or uneven clade representation (S12 Fig). Disparity during the Thanetian does exceed (Fig 3A, S13 Fig, S12 Table) that of the pre-extinction Campanian time-bin (PV_Campanian_=0.069; PV_Thanetian_=0.085; *p*=0.033), and was unaffected by pruning of synechodontiforms, a group of possible stem selachimorphs (Fig 3A and S13 Table). Comparisons with the Stevns Klint regional sub-sample found no significant change in tooth disparity (PV_lateMaastrichtian_=0.091; PV_earlyDanian_=0.069; *p*=0.106) across the K/Pg boundary event (Fig 3A and S14 Table). The extinction event was followed by a significant disparity increase between the early to middle Danian (PV_earlyDanian_=0.069; PV_middleDanian_=0.123; *p*=0.005).

**FIG. 3.**
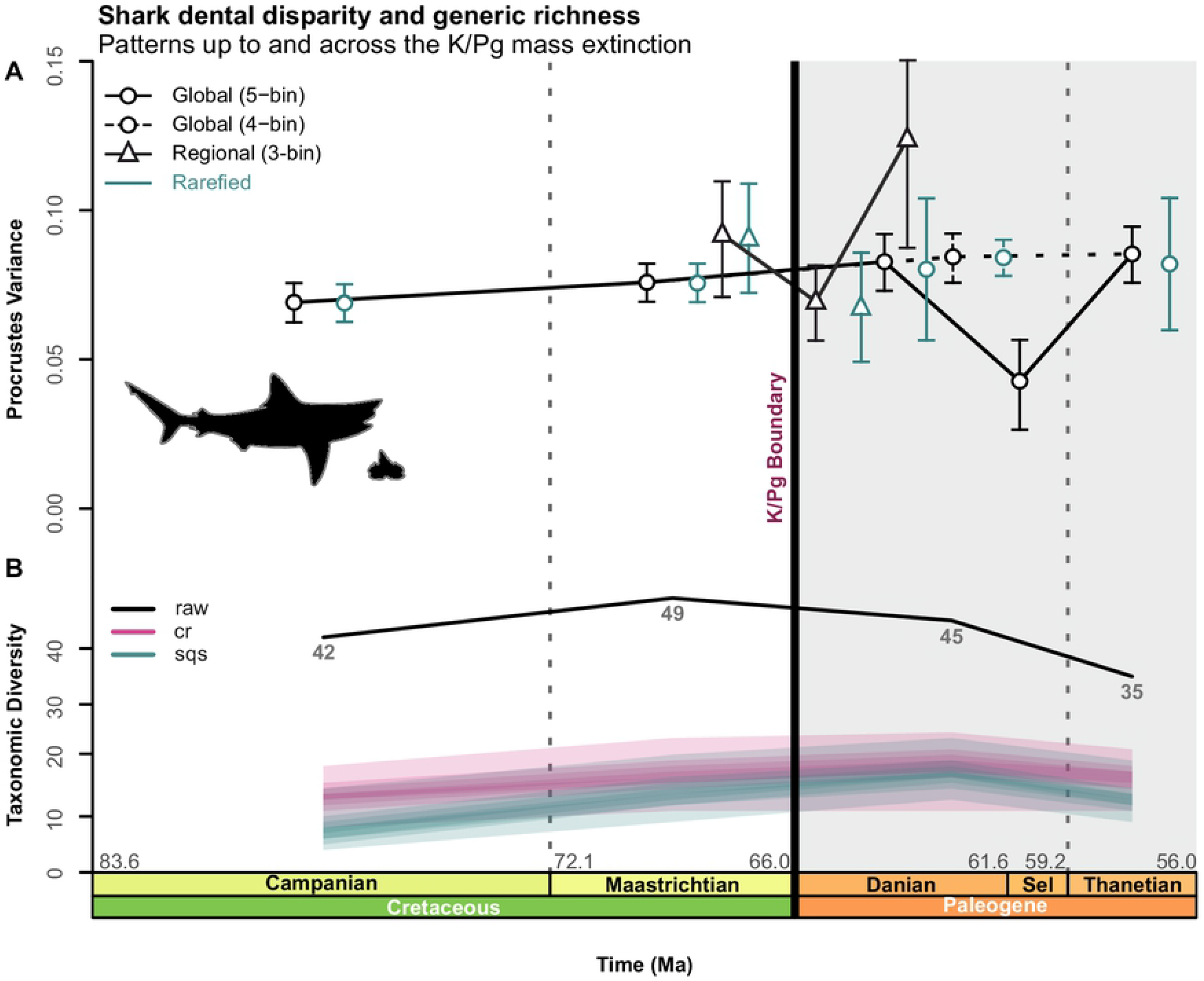
Global and regional level disparity of Selachimorpha. **(A)** Dental disparity with bootstrapped prediction intervals for the five and four-age time-binning schemes. The five-age time-binning scheme (n=1156) utilized sample-rarefaction on the lowest sampled Selandian time-bin (n=28). The four-age time-binning scheme (n=1198) used the Thanetian (n=187). Rarefied disparity is not shown for the Selandian (five-age time-bin) and middle Danian regional sub-sample (three-age time-bin). **(B)** Raw, rarefied, and SQS corrected taxonomic sampled-in-bin ‘SIB’ diversity through time. Shaded areas=quantiles. Sel=Selandian.

### Genus-level diversity

Raw genus-level occurrences showed a diversity increase in the Campanian to Maastrichtian time-bins, followed by a prolonged decline throughout the Danian–Thanetian (Fig 3B). However, sample-standardized curves using SQS and CR suggest stable diversity across the extinction event (Fig 3B).

### Superorder-level clade disparity

Relative stasis characterizes the disparity of both galeomorphs and squalomorphs across the Maastrichtian–Danian time-bins (Fig 4A–B, S15–17 Table). This result was consistent even after synechodontiforms were excluded (Fig 4A, S16 Table). Conversely, independent testing of the Stevns Klint regional sub-sample produced a significant disparity increase among galeomorphs (PV_earlyDanian_=0.069; PV_middleDanian_=0.109; *p*=0.022) in the early–middle Danian time-bins (Fig 4A, S18–19 Table), and a corresponding reduction in squalomorph disparity (Fig 4B, S20 Table) in the late Maastrichtian–early Danian time-bins (PV_lateMaastrichtian_=0.077; PV_earlyDanian_=0.041; *p*=0.005).

**FIG. 4.**
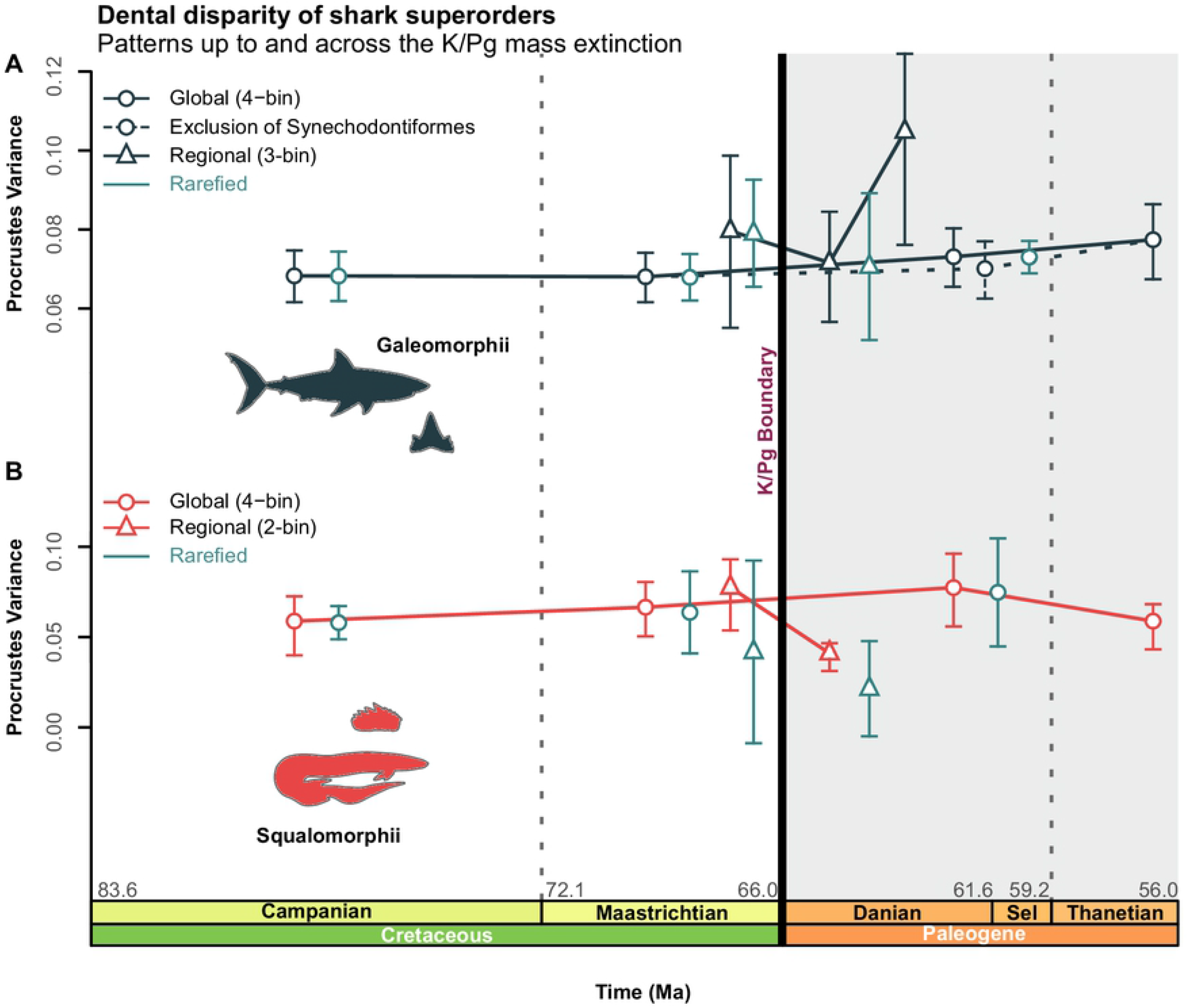
Superorder-level clade disparity profiles. **(A–B)** Global/regional raw and rarefied dental-disparity trajectories for Galeomorphii (including and excluding †Synechodontiformes) and Squalomorphii.

### Order-level clade disparity

Lamniform and carcharhiniform (Fig 5A–B, S21–22 Table) disparity returned maximum PV from the Campanian and Maastrichtian (Fig 5A–B). Global lamniform disparity is stable across the K/Pg boundary (PV_Maastrichtian_=0.070; PV_Danian-Selandian_=0.059; *p*=0.396; Fig 5A), and shows no significant change by the Thanetian (PV_Maastrichtian_=0.070; PV_Thanetian_=0.055; *p*=0.264; Fig 5A).

**FIG. 5.**
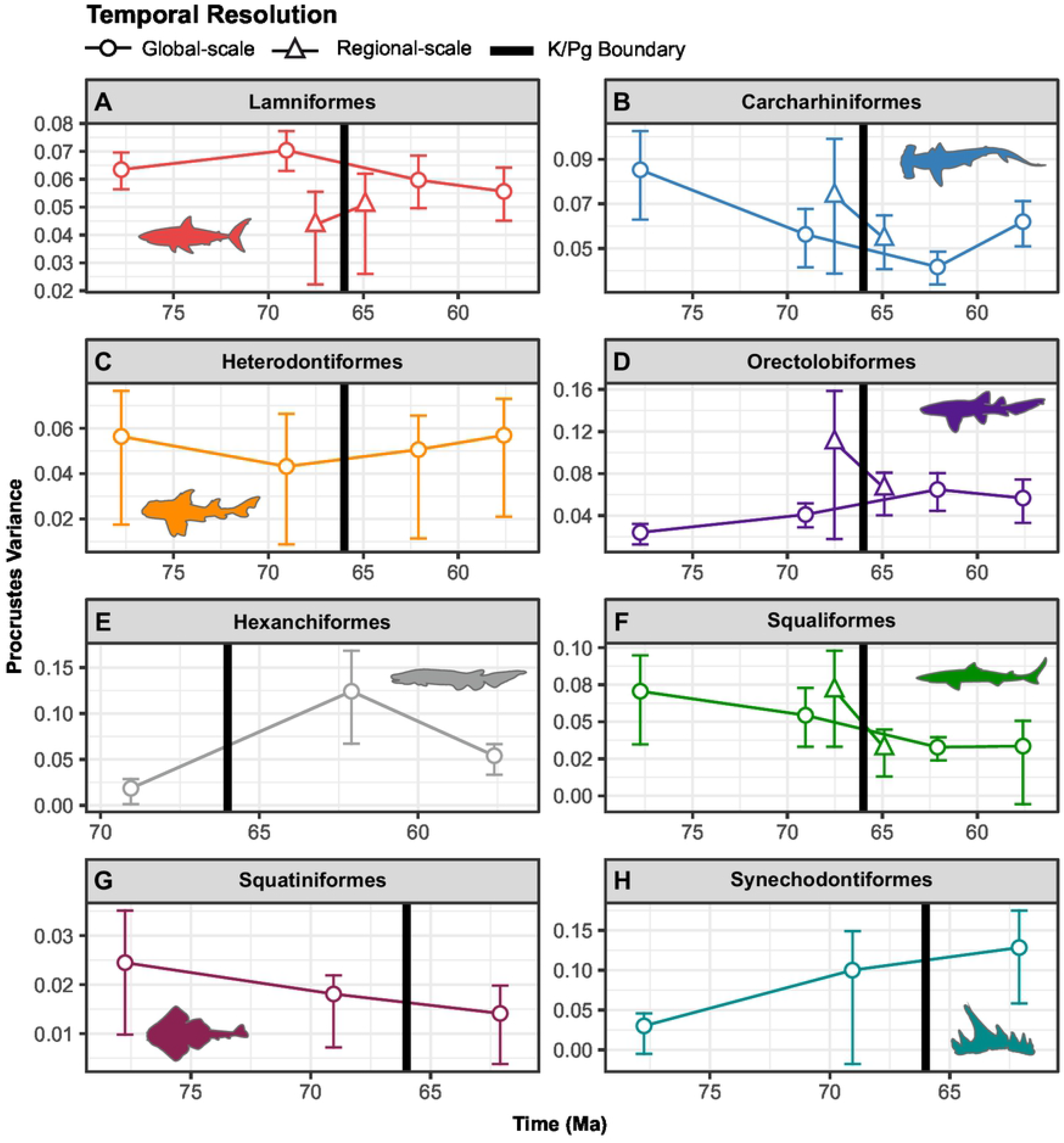
Global and regional clade-specific (order-level) disparity profiles. **(A–D)** Temporal resolution varies between groups but broadly tracks the time-binned K/Pg interval.

Carcharhiniforms exhibited a marked disparity decline from the Campanian to Danian-Selandian with our global sample, but not with the Stevns Klint regional sub-sample (Fig 5B, S22 Table). A distinct drop across the Campanian–Maastrichtian time-bins (PV_Campanian_=0.085; PV_Maastrichtian_=0.056; *p*=0.024) was followed by a non-significant change across the Maastrichtian–Danian-Selandian time-bins (PV_Maastrichtian_=0.056; PV_Danian-Selandian_=0.041; *p*=0.125), and a subsequent disparity increase from the Danian-Selandian– Thanetian (PV_Danian–Selandian_=0.041; PV_Thanetian_=0.061; *p*=0.020) (Fig. 5B).

Heterodontiform disparity was uniform across the Campanian to Thanetian time-bins (Fig 5C, S23 Table). On the other hand, orectolobiforms showed a significant disparity increase (PV_Campanian_=0.024; PV_Danian–Selandian_=0.064; *p*=0.016) from the Campanian to Danian-Selandian time-bins (Fig 5D, S24 Table). Conversely, we recovered no significant change in orectolobiform disparity between the Maastrichtian–Danian-Selandian time-bins (PV_Maastrichtian_=0.041; PV_Danian–Selandian_=0.064; *p*=0.096), and neither while using our regional sub-sample (PV_lateMaastrichtian_=0.109; PV_earlyDanian_= 0.066; *p*=0.294) (Fig. 5D).

Hexanchiforms disparity was statistically non-differentiable across the Maastrichtian–Danian-Selandian time-bins (PV_Maastrichtian_=0.018; PV_Danian–Selandian_=0.124; *p*=0.104), and also between the Danian-Selandian and Thanetian time-bins (PV_Danian–Selandian_ =0.124; PV_Thanetian_=0.054; *p*=0.072) (Fig 5E, S25 Table). Squaliforms similarly showed stable disparity from the Maastrichtian–Danian-Selandian time-bins (*p*=0.084), both with global, and Stevns Klint regional sub-sample data (Fig 5F, S26 Table). Finally, squatiniforms and synechodontiforms were stable, but their small sample sizes make interpretations equivocal (Fig 5G–H, S27–S28 Table).

### Global partial disparity

Our PV values indicate that selachimorph global disparity during the Campanian and Maastrichtian was predominantly driven by lamniforms (Fig 6A). Not surprisingly, lamniforms were also the most numerically abundant taxa in our dataset (S1 Fig). Lamniform global disparity subsequently decreased in the Danian time-bin (Fig 6A), but increased again in the Selandian time-bin. Altogether, lamniforms accounted for almost the total global variance observed within Selachimorpha (Fig 6A). Carcharhiniforms and hexanchiforms otherwise contributed towards a global disparity increase across the Danian–Thanetian time-bins (Fig 6A).

**FIG. 6.**
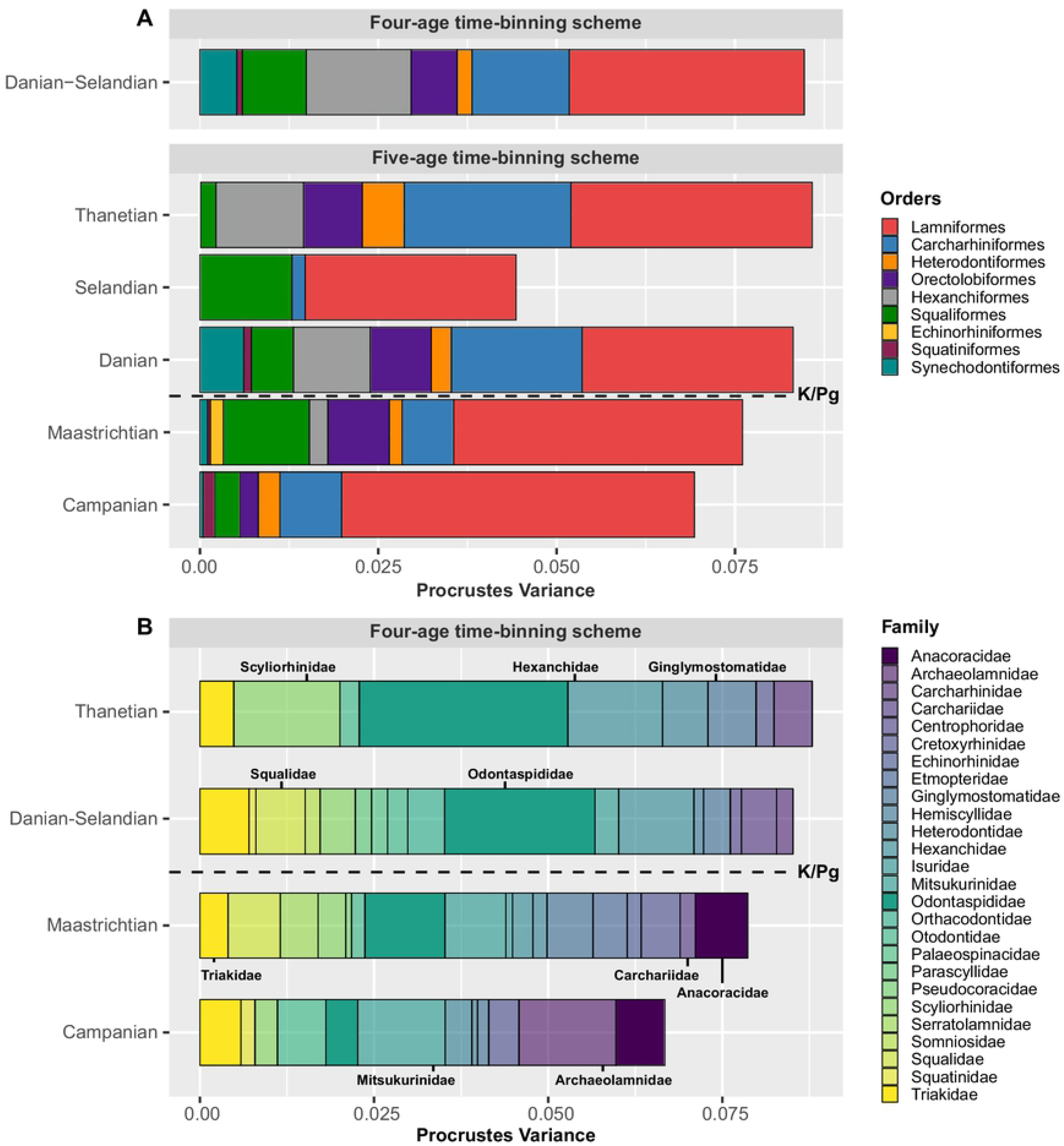
Order and family level partial disparity. **(A–B)** Grouped bar plot depicting clade-level contributions (pPV) to overall disparity in each time-bin. **(B)** Families with sample sizes of n<5 were omitted. Resulting data: a total of 26 families and 1039 specimens when using the global four-age time-binning scheme.

Although sampling is admittedly uneven, our break-down of family-level clade disparity (Fig. 6B) indicates that archaeolamnids, mitsukurinids, and anacoracids are the primary drivers within the Campanian time-bin. Odontaspidid sharks increase their disparity following the extinction of anacoracids in the Danian-Selandian time-bin, and expanded substantially in the Thanetian time-bin (Fig. 6B). In correspondence, triakids, hexanchids, and squalids were the major non-lamniform contributors to disparity in the Danian-Selandian time-bin, with scyliorhinids becoming dominant in the Thanetian time-bin. Other family-level clades, such as ginglymostomatids and hexanchids also increased their disparity from the Danian-Selandian to Thanetian time-bins (Fig. 6B).

### Geographic distribution of disparity

Samples from the Western Interior Basin (58% derived entirely from the upper Judith River Formation in the U.S.A) of North America and the Central and Northern European basins accounted for most of the disparity in our Campanian time-bin (Fig 7A and S14 Fig). Stratigraphically equivalent sequences from the Ouled Abdoun and Tarfaya basins of Morocco (N=3) contributed comparatively limited disparity, although, the Moroccan sample contributes substantially to the Maastrichtian signal (Fig 7B). In general, geographic regions represented by small sub-samples exhibited low disparities, unlike large sub-sampled regions, such as the U.S.A., which overwhelmingly contributed to global disparity in our Maastrichtian time-bin (Fig 7B). Sub-age comparisons indicated that the U.S.A. sub-sample was less dominant in the Maastrichtian time-bin, and was outweighed by the Stevns Klint xsub-sample combined with the Limhamn Quarry sub-sample from Sweden in our Danian– Selandian time-bins (Fig 7C). Both the U.S.A. and Morocco sub-samples returned to high disparity in the Thanetian time-bin (Fig 7D).

**FIG. 7.**
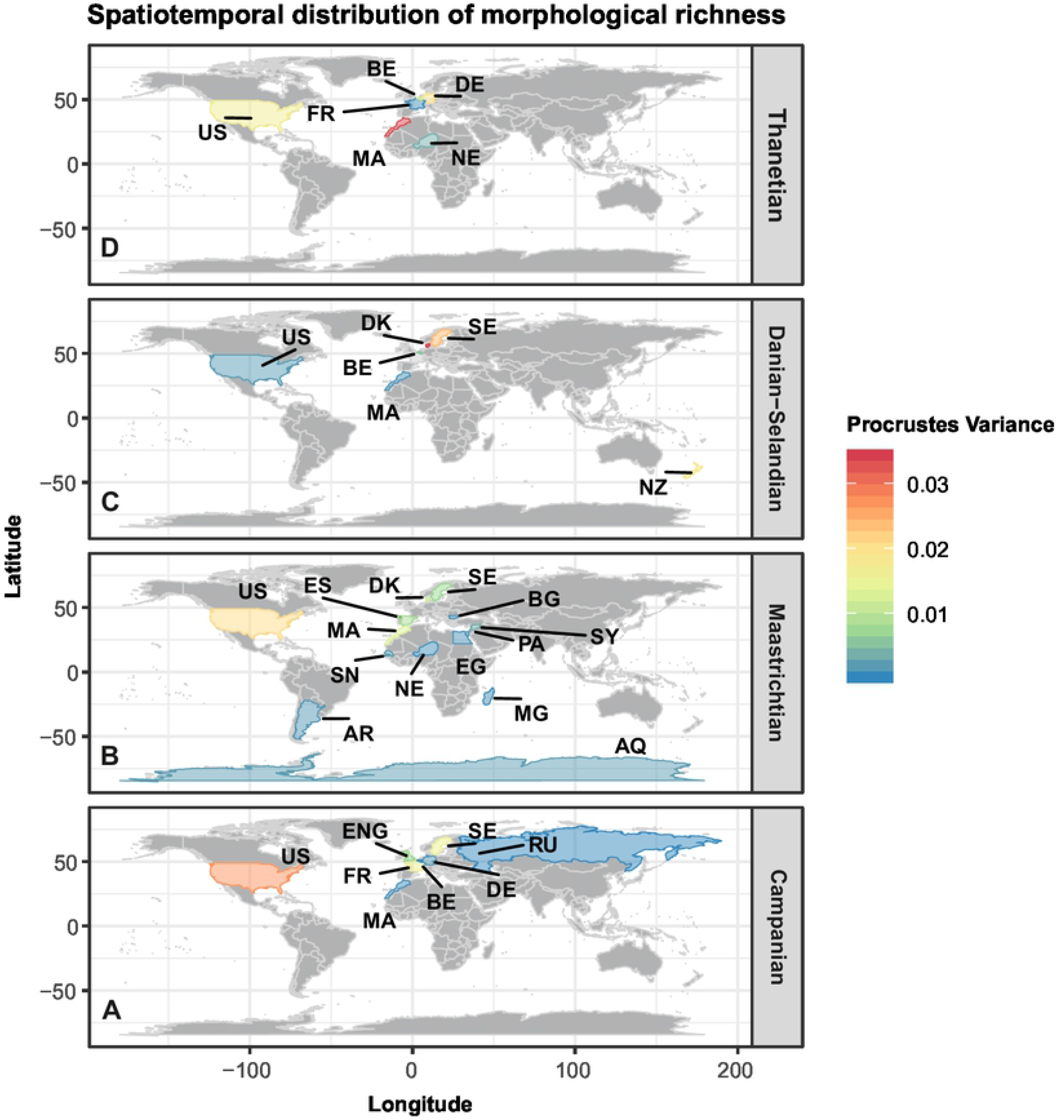
Geographic distribution of disparity. **(A–D)** Partial disparities mapped using the global four-age time-binning scheme. Colour gradient represents low-to-high disparity.

### Effects of heterodonty on disparity

A significant disparity increase occurred across the Maastrichtian–Danian-Selandian time-bins (PV_Maastrichtian_=0.066; PV_Danian–Selandian=_ 0.082; *p*=0.032) with our monognathic heterodonty model (S29–S30 Table). The combined monognathic and dignathic heterodonty model also produced a disparity increase from the Campanian to Maastrichtian time-bins (*p*=0.016). However, dignathic heterodonty, together with a multivariate monognathic/dignathic heterodonty interaction model yielded no comparable disparity shifts (S15 Fig and S29–S30 Table) across the K/Pg boundary. Finally, model comparisons revealed largely consistent trajectories in dental disparity throughout the Campanian–Thanetian interval, despite some effect (i.e., the inflation of absolute values) of heterodonty (S15 Fig).

### Global morphospace

Pairwise comparisons along all PCs found significant differences in morphospace distribution between time-bins (Fig 8, Table 2). The PC1 distributions are platykurtic (kurtosis < 3) and positively skewed, except during the Thanetian, which was negatively skewed (*g*1_Thanetian_= − 0.0336). Computed distribution-specific interquartile ranges (IQR) show minimal morphospace dispersion on PC2 (S31–S32 Table). A Hartigan’s dip test of multimodality did not reject a unimodal distribution on either PC1 or PC2. However, a significant positive morphospace shift occurred between the Maastrichtian–Danian along PC1 (Fig 8A, S16 Fig, S31–S32 Table). Notably, there was no corresponding change in the modal-shape configurations, although a reduction in positively loaded morphospace accompanied range-shortening among the minimum values (Fig 8A, S16 Fig).

**Table 2.** Non-parametric analysis of variance based on RRPP for all PCs. Coefficient estimation via ordinary least squares (OLS). Type I (sequential) sums of squares were used to calculate sums of squares and cross-products matrices. Effect sizes (Z) based on F distribution.

|  | *d.f.* | SS | MS | R<sup>2</sup> | F | Z | Pr(>F) |
| --- | --- | --- | --- | --- | --- | --- | --- |
| **Selachimorpha** |  |  |  |  |  |  |  |
| ~ interaction (global 5-bin) | 4 | 2.625 | 0.65621 | 0.02911 | 8.6269 | 5.0104 | **0.001\*\*** |
| ~ interaction (global 4-bin) | 3 | 2.219 | 0.73957 | 0.0234 | 9.5354 | 4.9451 | **0.001\*\*** |
| ~ interaction (regional 3-bin) | 2 | 0.840 | 0.41999 | 0.06043 | 4.8235 | 2.8145 | **0.002\*\*** |
| **Galeomorphii**<br>**(Synechodontiformes included)** |  |  |  |  |  |  |  |
| ~ interaction (global 4-bin) | 3 | 2.254 | 0.75136 | 0.02966 | 10.587 | 5.0617 | **0.001\*\*** |
| ~ interaction (regional 3-bin) | 2 | 0.7811 | 0.39053 | 0.07727 | 4.7316 | 2.796 | **0.003\*\*** |
| **Squalomorphii** |  |  |  |  |  |  |  |
| ~ interaction (global 4-bin) | 3 | 0.9088 | 0.302947 | 0.07875 | 4.3028 | 3.8612 | **0.001\*\*** |
| ~ interaction (regional 3-bin) | 2 | 0.30381 | 0.151907 | 0.12846 | 2.5058 | 1.9687 | **0.015\*** |

Symbols: *d.f.* = degrees of freedom; SS = sequential sums of squares; MS = Mean squares; F statistics = F value by permutation; R<sup>2</sup> = coefficient of determination; Z = effect size. *P*-values are based on 999 permutations.

**FIG. 8.**
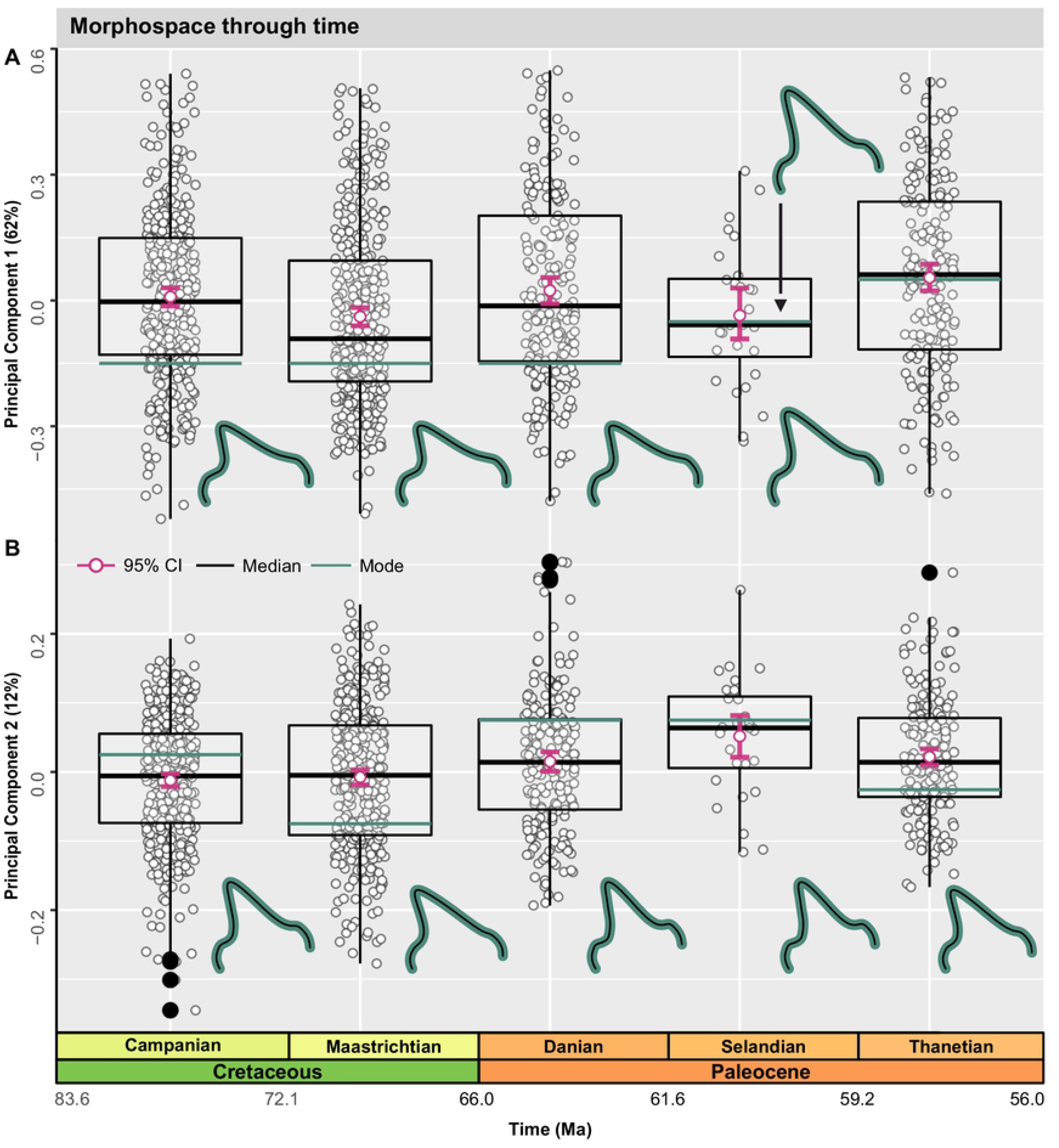
Global morphospace time-series. **(A-B)** Jittered box-plots visualizing the distribution of time-bins along PC1 and PC2. Graph depicts patterns of overall shape change using the five-age time-binning scheme. Arithmetic mean with 95% confidence limits were derived via non-parametric bootstrapping. Modal and median values are shown along PC1 and PC2; TPS-grids indicate changes in modal value between time-bins.

The average value along PC1 shifted positively from the Selandian to the Thanetian (Fig 8A). Comparisons between the Campanian–Maastrichtian (*p*=0.025) and Maastrichtian versus Thanetian time-bins (*p*=0.013) likewise yielded significant morphospace shifts (S33 Table).

The PC2 distribution was leptokurtic (high kurtosis > 3) during the Campanian and Danian, but characterized by low kurtosis from the other time-bins (Fig 8B). A positive shift occurred from the Maastrichtian (*g*1_Maastrichtian_= −0.0806) to the Danian (*g*1_Danian_=0.2833) (Fig 8B, S31 Table), and coincide with a loss of negatively loaded tooth morphologies (Fig 8B).

Nonetheless, the positive loading frequency diminished during the Danian, and a Procrustes ANOVA also found statistical differentiation between the Campanian–Danian (*p*=0.008) and Maastrichtian–Danian (*p*=0.015) time-bins (S33 Table). A comparable morphospace shift across the K/Pg interval was detected with the four-age time-binning scheme (S13 Fig), and incorporated subtle changes in modal-shape within the Campanian + Maastrichtian–Danian-Selandian time-bins (S16 Fig).

### Regional morphospace

The Stevns Klint regional sub-sample reveals no substantial shifts along PC1 or PC2 from the late Maastrichtian to the early Danian (Fig 9A–B, S33 Table). However, a significant sub-age positive shift in mean morphology occurred along PC1 between the late Maastrichtian–middle Danian (*p*=0.005), and also between the early–middle Danian time-bins (*p*=0.005) (Fig 9A, S33 Table). Both, the late Maastrichtian (*g*1_late Maastrichtian_=0.2640) and early Danian time-bins (*g*1_early Danian_=0.2957) are characterized by positively skewed distributions, but with negative skewing during the middle Danian (*g*1_middle Danian_= −0.1614). An increase in negatively loaded teeth is associated with the early Danian along PC1; a pattern also reflected in the modal-shape configuration (Fig. 9A).

**FIG. 9.**
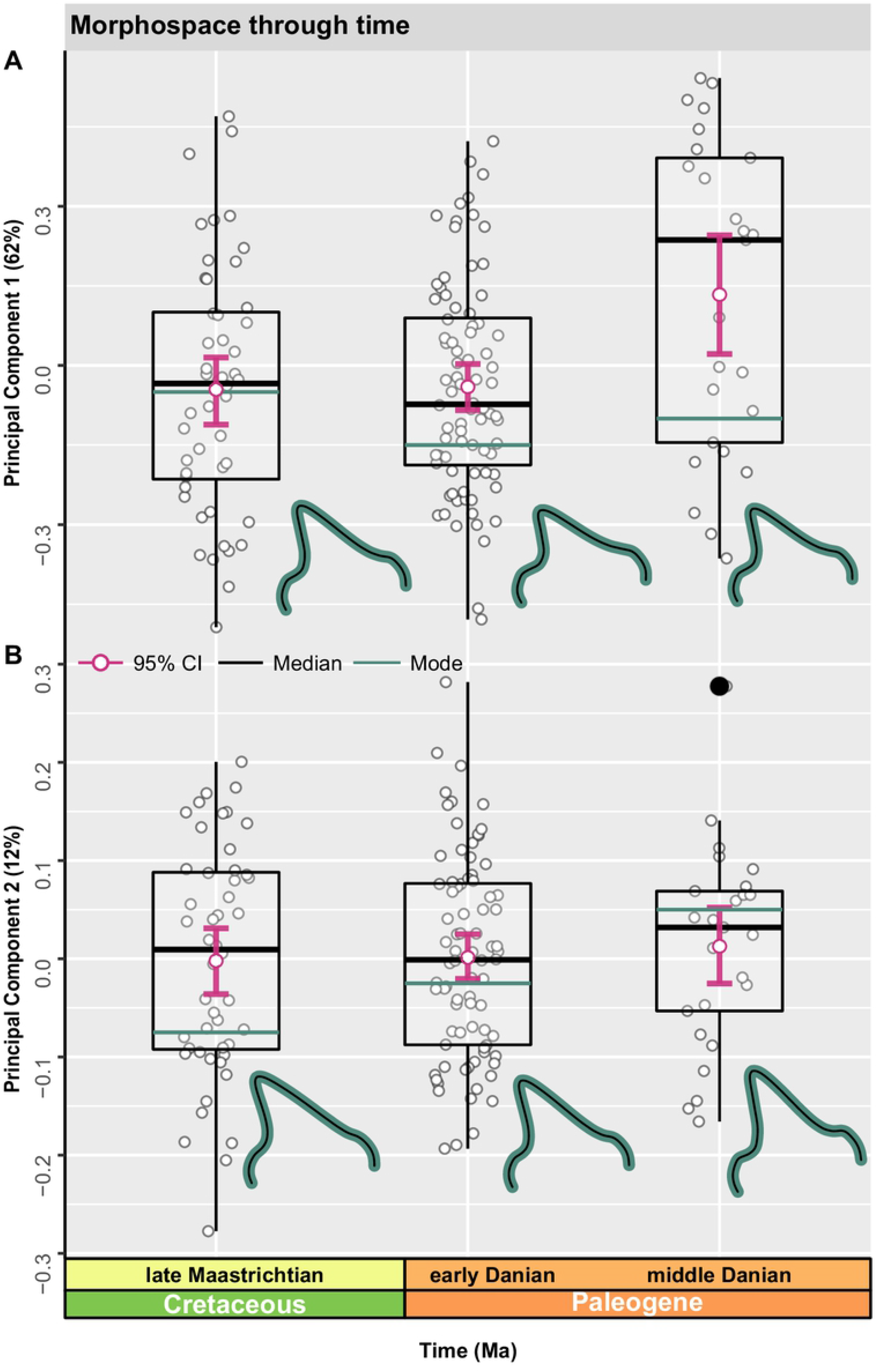
Regional morphospace time-series. **(A–B)** Jittered box-plots visualizing the distribution of time-bins along PC1 and PC2 (n=153). TPS-grids correspond to modal values within each time-bin.

The PC2 distribution was negatively skewed in the late Maastrichtian time-bin, while the early–middle Danian time-bins exhibited positive skewing (Fig 9B, S33 Table). Modal shape changes were more pronounced along PC2 (Fig 9B), with a gradual shift from negatively to positively loaded values (Fig 9B).

### Superorder-level clade morphospace

Galeomorphs and squalomorphs occupied comparable regions on PC1 throughout the entire Maastrichtian–Thanetian interval (Fig 10A), but exhibit notable differences in their mean and medial values. On average, galeomorphs are characterised by tall, narrow teeth, whereas squalomorphs are low crowned. The most pronounced shift occurs across the K/Pg Boundary along PC1, as well as on PC3 and PC4 (S34 Table), where galeomorphs exhibit a reduction in negative PC1 values (Fig 10A). Squalomorphs underwent a significant positive shift on PC2 (*p*=0.016) from the Maastrichtian to the Danian-Selandian (Fig 10A–B, S35 Table).

**FIG. 10.**
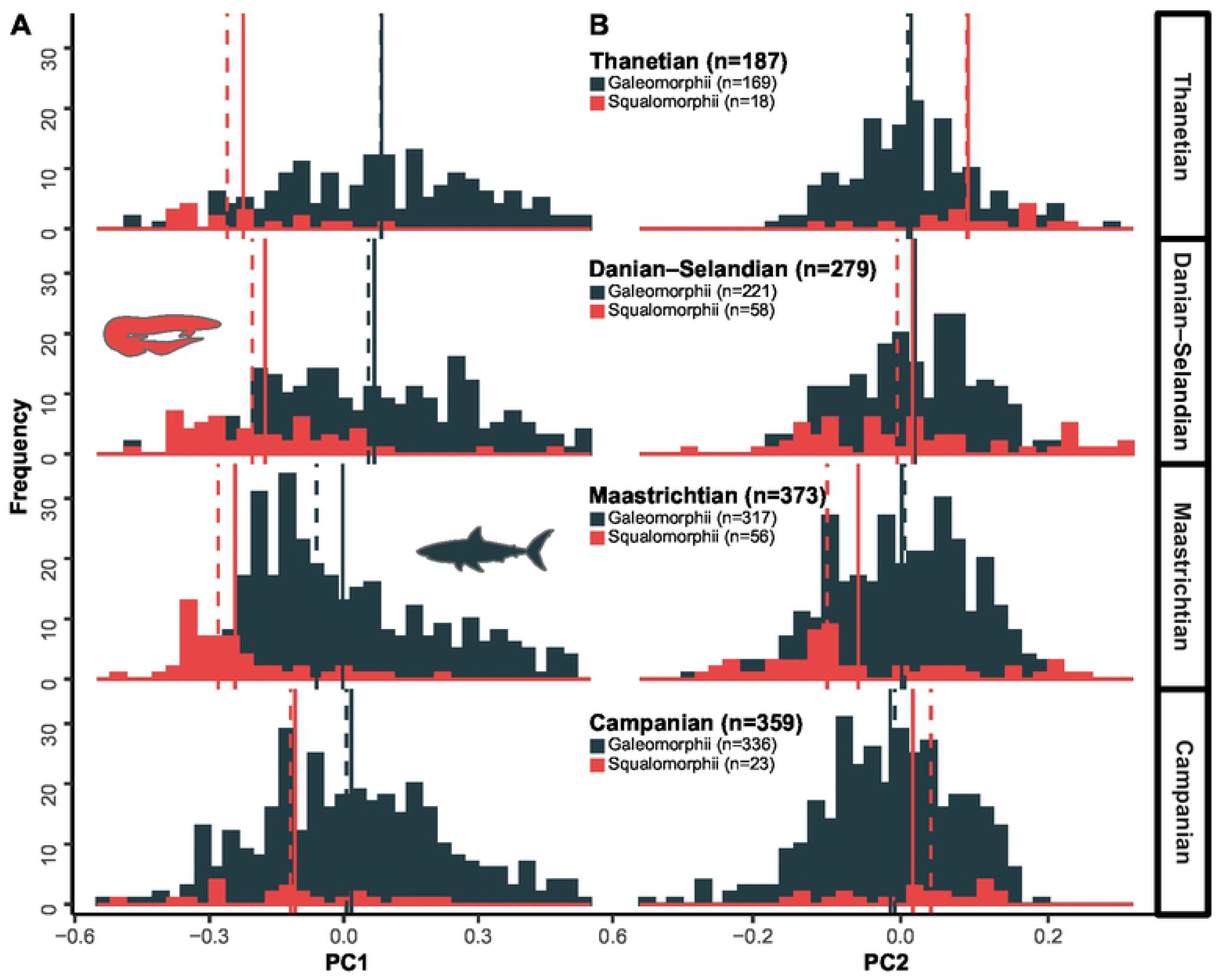
Superorder-level clade morphospace using the global four-age time-binning scheme. **(A–D)** Frequency histogram plots for PC1–PC2. Measures of tendency, including the median (dashed line) and arithmetic (solid line) mean are shown. Clade sample sizes by time-bin are indicated.

### Order-level clade morphospace

Lamniforms display a significant positive shift in mean morphology along PC1 from the Maastrichtian to Danian-Selandian time-bins (*p*=0.002) (Fig 11A, S36 Table), concomitant with a shift in distribution patterns from positively to negatively skewed (*g*1_Maastrichitan_=0.5035; *g*1_Danian–Selandian_=–0.3193). The frequency of negatively loaded morphologies on PC1 is low during the Thanetian; the distribution of which is significantly different from both the Campanian (*p*=0.002) and the Maastrichtian (*p*=0.002).

**FIG. 11.**
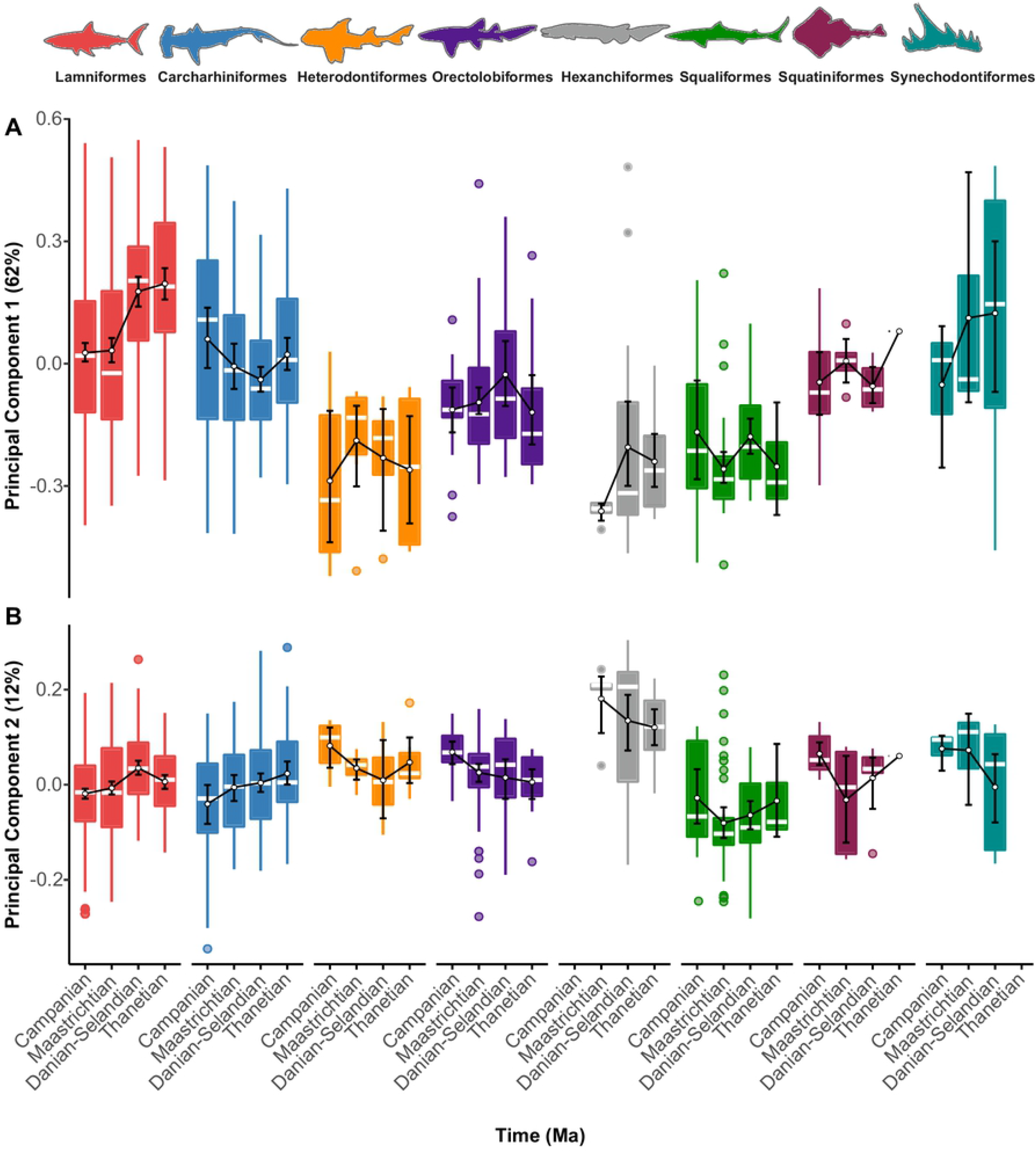
Order-level clade morphospace using the four-age time-binning scheme. **(A–B)** Patterns along PC1 and PC2. **(C–D)** Minimum and maximum values along the ordinated axes. Sampling was insufficient to estimate reliable disparity for Echinorhiniformes. Measures of tendency, including the median and arithmetic mean are shown with computed 95% non-parametric bootstrap confidence intervals and potential outliers.

Carcharhiniforms alternatively exhibited a relatively stable morphospace along PC1 across the K/Pg Boundary, although their distribution contracted (IQR_Maastrichitan_=0.25 vs. IQR_Danian– Selandian_=0.19). A significant negative shift between the Campanian and Danian–Selandian time-bins (*p*=0.024) is noted, along with a possible gradual transition from generally positive to negative values (S37 Table), via the contraction of positive PC1 values (Fig 11A). Heterodontiforms, orectolobiforms, hexanchiforms, squaliforms, squatiniforms, and synechodontiforms all produced no significant deviations along PC1 in the Campanian to Thanetian time-bins (Fig 11A, S38–S43Table).

On PC2, lamniforms exhibited a depletion of negative values from the Maastrichtian–Danian-Selandian time-bins, and these are statistically different (*p*=0.004) (Fig 11B). Pairwise comparisons also recovered a significant difference between the Campanian and Danian– Selandian time-bins (*p*=0.004). By contrast, carcharhiniforms show no major changes along PC2, except for a slight increase in positive skewness across the Danian-Selandian–Thanetian time-bins, which might imply an exploration of new morphospace (Fig 11B). We additionally detected significant differences between the Campanian and Danian–Selandian (*p*=0.031), and the Campanian and Thanetian time-bins (*p*=0.007), although this signal reduced after FDR-adjustment of the *p*-values (S37 Table). No notable shifts were noted along PC2 for heterodontiforms and orectolobiforms (Fig 11B). Yet, these clades did incline towards positive skewness. Finally, no significant shifts in average morphology were noted along PC2 among squaliforms, squatiniforms, and synechodontiforms (S41–43 Table).

## DISCUSSION

### Disparity dynamics across the K/Pg Boundary

Our results indicate that the global disparity of selachimorphs across the K/Pg extinction event was largely stable. Stasis is consistent with previous dental disparity-based studies of sharks [20] and interpretations of the elasmobranch fossil record [16], but contrasts with significant losses in species richness of almost 50% [17,18,77], suggesting shark diversity and disparity were decoupled across the K/Pg Boundary. Importantly, our evidence for static disparity in selachimorphs across the boundary is retained after accounting for heterodonty and differences in order-level representation between the K/Pg time-bins (S15 Fig). Overall, we find that these sources of variation only alter the disparity inferred for the Thanetian that, upon correction, cannot be differentiated from that of the Campanian (see Fig. 3A, S15 Fig, and S29 Table). In either case, morphological stasis about the K/Pg Boundary suggests that the loss of 17% of families [17] was not linked to major losses in ecological diversity and, overall, conforms to a ‘non-selective’ extinction model [78]. However, at lower taxonomic scales, lamniforms experienced the selective extinction of Cretaceous anacoracids [20] with triangular, blade-like teeth (Fig 11A–B) and, in the aftermath, lamniform morphospace underwent a noticeable Paleocene proliferation of apicobasally tall, laterally cusped teeth, driven by odontaspidids (Fig 11A–B). This dynamic of loss in one area of morphospace and subsequent proliferation in another supports a ‘shift’ extinction model [20], the result of which is a stable disparity. Concomitantly, carcharhiniforms characterized by mesiodistally broad and low teeth proliferated during the Paleocene (Fig 11A–B), perhaps reflecting a ‘non-adaptive’ radiation along with minimal ecological divergence [20]. In comparison to galeomorphs, squalomorph disparity seemed largely unaffected by the extinction event, despite previously identified moderate taxonomic losses among squalids [17]. Much like the overall pattern, squalids may have undergone a non-selective extinction. However, squalomorphs remain poorly sampled across the end-Cretaceous extinction event (S1 Table), and associated patterns should be interpreted cautiously.

Despite the proliferation of certain tooth types in the aftermath of the extinction event, static disparity does not support a Paleocene diversification of sharks, which is in contrast to the substantial ecological and taxic diversification of coeval actinopterygian fishes (e.g., acanthomorphs and carangarians) [16,19,79–82]. Nevertheless, Sibert et al. [83] recently found that post-Mesozoic actinopterygian disparity underwent alteration in only a few dominant tooth morphotypes. Likewise, we suggest that while, in general, sharks do not seem to have experienced much change in disparity across the K/Pg boundary, they did suffer sufficient ecological disturbance to trigger a compositional transformation in the diversity of certain constituent clades [20,83], such as lamniform and carcharhiniforms.

### Regional-and global-level extinction dynamics

Adolfssen & Ward [64] reported a 33% decline in chondrichthyan richness across the K/Pg extinction interval sampled at Stevns Klint in Denmark, which is substantially less than that reported from Morocco (∼96%: [84]) or globally (∼84%: [17]). Stevns Klint preserves a largely endemic Boreal fauna [64], yet its static disparity signal is largely indistinguishable from that recovered by our global disparity analysis. However, a slight disparity increase (Fig. 3A) was recovered from the early to middle Danian, which conforms to an overall increase (albeit statistically non-significant) in absolute values amongst galeomorphs (Fig. 4A). This regional pattern is indicative of likely short-term recovery in the aftermath of the extinction event, but otherwise posit a direct correlation with distribution trends evidenced globally for the early Paleocene.

On a broader level, regional disparity occurred in ‘hotspots’ that transitioned over the sampled interval. Results indicate that (1) the Western Interior Basin assemblages from the U.S.A. contributed to much of the measurable global disparity during the Campanian and Maastrichtian (Fig 7A–B); (2) the Scandinavian Boreal assemblages from Denmark and Sweden likewise predominate disparity for the K/Pg extinction interval (Fig 7C and S14 Fig); and (3) the Mediterranean Tethyan and Atlantic margin basins of Morocco provide most of the disparity for the Thanetian (Fig 7D). While these results are clearly influenced by preservation biases (e.g., spatial variation in sampling across the Cretaceous/Paleogene Boundary) (as is typical of many fossil vertebrates: [85]), they also highlight underlying provincialism, and noticeably correlate with the changing depositional context of northern mid-latitude epicontinental habitats during the Cretaceous–Paleocene timeframe. For instance, the steady disparity decline in North America correlates with the regression of the Western Interior Seaway across this interval [86]. Therefore, despite overall consistent patterns between global and regional disparity profiles (when assessing the Denmark sub-sample), we suggest that extinction and recovery dynamics of sharks varied globally, as previously documented for other post-mass extinction marine ecosystems [87–89]. Moreover, it is possibly that selachimorphs survived the K/Pg crisis in epeiric refugia [64] and subsequently ‘radiated’ from localised diversification centers.

### Ecological implications

The reorganization of selachimorph communities across the K/Pg mass extinction event [17,18,20,24] can almost exclusively be attributed to large body-sized, apex-predatory anacoracid lamniforms (e.g., the reduction of negatively loaded tooth structures) [20,21]. Such patterns are consistent with the loss of other marine apex predators, such as mosasaurs and plesiosaurs at the end of the Mesozoic [90–92]. Unlike marine tetrapods, which would not return until the Eocene, the extinction of Cretaceous anacoracids was immediately followed by the appearance of large-bodied hexanchiforms in some earliest Paleocene deposits (e.g., the Limhamn quarry) [93]. Anacoracids and hexanchids exhibit substantial morphospace overlap, especially along PC1, (S17 Fig), suggesting at least some ecological equivalency, which is supported by modern-day observations that the hexanchid *Notorynchus cepedianus* becomes more abundant following the removal of the large-bodied lamniform, *Carcharodon carcharias* [94]. Unfortunately, there is at present no universal approach for estimating the body sizes of fossil sharks from their teeth (but see: [95]), and further paleoecological comparisons between hexanchids and lamniforms remain ambiguous. Regardless, the continuity of large-bodied sharks across the boundary, does suggest that extinction susceptibility was not limited to relative body-size. Indeed, studies on living cartilaginous fishes have shown that habitat preferences and reproductive strategies play a more important role in determining survival, whereas, body-size has a modest effect on extinction risk [96].

Finally, for sharks, the K/Pg event should also be viewed in terms of diversification, rather than simply extinction. In particular, lamniforms (odontaspidids) and carcharhiniforms (triakids, scyliorhinids) morphologically proliferated in the early Paleocene, potentially taking advantage of teleost fishes as an newly emergent food resource [16]. These patterns, highlight the importance of biotic factors (i.e., predator-prey relationship) within the context of global mass extinction, and it’s contrasting effect among shark groups.

## CONCLUSIONS

Understanding the evolutionary dynamics of sharks during the catastrophic end-Cretaceous mass extinction event [17,18,20,21,77] has analytically lagged behind assessments of other dominant marine vertebrate groups such as teleost fishes [16,19,79–82], and marine/freshwater reptiles [91,92,97,98]. Consequently, we present the first comprehensive geometric morphometric evaluation of selachimorph disparity across multiple clades based on their prolific dental fossil record. The combined observation of virtually static levels of morphological disparity and the potential for differential extinction dynamics across the globe suggest that, in a broad sense, the K/Pg extinction event was not as severe for sharks, as it was for other vertebrates [90,91,99,100] Nevertheless, one particularly dominant Mesozoic clade of sharks, anacoracid lamniforms, underwent a selective extinction, which saw to the loss of triangular, blade-like teeth that were perhaps associated with feeding on large-bodied contemporaneous tetrapods [20]. Other coeval selachimorph clades, however, survived and appear of have been largely unaffected by the extinction event, with some groups of carcharhiniforms and lamniforms even proliferating during the Paleocene. We interpret this as an extinction-mediated ecological ‘shift’, whereby certain sharks underwent significant changes in their morphology, but without substantial modifications to their overall disparity [20]. The potential drivers of this transition include possible shifts in food resources, supported by the Paleocene diversification of teleost fishes, although other mechanisms cannot be rejected at this time. Finally, our application of a morphospace-disparity framework offers a powerful interpretive tool that compliments traditional taxonomy-based assessments of discrete character data [e.g., 101,102]. We therefore advocate similar continuously valued disparity descriptors to reconstruct deep time evolutionary patterns in the fossil record of sharks in the future, while acknowledging that variation sources, such as heterodonty and differential spatiotemporal sampling, need to be accommodated.

## ACKNOWLEDGEMENTS

We thank Jesper Milan (Geomuseum Faxe) and Mikael Siversson (Western Australian Museum) for discussions and access to material. Daniel Snitting (Uppsala University) assisted with processing of image data. Julien Claude (Université de Montpellier), Göran Arnqvist (Uppsala University), Grahame Lloyd (University of Leeds), Dean Adams (Iowa State University), and Michael L. Collyer (Chatham University) contributed expertise on the morphometric and statistical analyses. Our research was financed by The Royal Swedish Academy of Sciences (GS2017-0018) to M.B., and a Wallenberg Scholarship from the Knut and Alice Wallenberg Foundation to P.E.A. B.P.K. also acknowledges funding from a Swedish Research Council Project Grant (2020-3423), and N.E.C. is funded by an Australian Research Council Discovery Early Career Research Grant (DE190101423).

## DATA ARCHIVING STATEMENT

Data and supporting information for this study are available in the Dryad Digital Repository.

